# Finetuning Foundation Models for Temporal Clinical Transcriptomics Data

**DOI:** 10.1101/2025.10.20.683402

**Authors:** Sachin Mathur, Alexander Kagan, Peyman Passban, Hamid Mattoo, Euxhen Hasanaj, Ziv Bar-Joseph

**Author notes:** Contributing authors.

## Abstract

**Background:** Timeseries clinical transcriptomic datasets offer the opportunity to gain insights into the dynamics of disease mechanisms/treatment responses. However, their utility in uncovering temporal patterns is often limited by high noise levels and small sample sizes. Leveraging foundational gene embeddings and incorporating interaction information can help address these challenges, improve gene network analysis, and enable the detection of subtle changes that drive disease progression or drug response.

**Results:** We finetuned gene embeddings from foundation models using healthy tissue gene expression data and used them in temporal GNNs to model gene expression of responder and non-responders to treatment in 3 disease datasets - ulcerative colitis, Crohn’s disease and psoriasis. Application of our method to these datasets confirmed known mechanisms associated with drug action, and also identified key differences between activated and repressed pathways for responders and non responders including B-Cell activation and mitochondria related activity in ulcerative colitis patients.

**Conclusion:** Finetuning gene embeddings from foundation models provide a richer context to model gene expression data compared to using them in their naive state. Even with smaller sample sizes, results from GNN-based temporal models outperform traditional methods by detecting known mechanisms of response and unraveling role of genes and mechanisms not known to be associated with response and non-response.

**Code Availability:** Code and data are available in a public GitHub repository - https://github.com/Sanofi-Public/GNN-Timeseries

## 1 Introduction

Timeseries clinical transcriptomic datasets allow understanding of dynamics of disease progression/treatment response in patients and help to identify drug targets and discover disease endotypes. In most clinical trials or observational studies, patients show variable response to drug treatment (Boessen et al. 2012). While some patients benefit from the drug (responder group), others show limited to no response (non-responder group). This could be due to various reasons such as differences in genetic characteristics, presence of distinct disease subtypes, environmental factors or differences in disease severity. It is important to understand the reasons for response/non-response at the molecular level so that new drug targets can be found to benefit patients who do not respond to standard therapies, which is very challenging. In addition to the high resolution of the expression data and the small number of individuals that are profiled, which often makes it hard to correctly identify an underlying cause, clinical transcriptomic datasets tend to be noisy because of heterogeneity in patient demographics, disease state, environmental factors, etc.

Several methods have been used to analyze and model clinical transcriptomic data. (Oh and Li 2021) provides a comprehensive review of different timeseries methods for bulk RNASeq datasets. Traditional methods such as Dynamic Bayesian networks (DBNs) (Ajmal and Madden 2021) have been widely used to model temporal dependencies to identify causal connections, but they fail to fully account for the topological structure of biological networks. DBNs require substantial computational resources when analyzing more than 1500 genes, which necessitates placing stringent filtering criteria and risks removing genes of interest. MASigPro (Conesa et al. 2006) uses a regression based approach to find gene expression profile differences, while TiSA (Lefol et al. 2023) combines differential gene expression and clustering based on recursive thresholding to detect differential gene expression. Newer methods such as constrained pseudo-ordering of samples (Mathur et al. 2024), and PhenoPath (Campbell and Yau 2018) adopt different approaches to address variable disease states of patients and noise issues by ordering samples on a pseudo-disease axis. However, these methods do not utilize prior knowledge such as protein-protein interaction or gene regulatory networks in their implementation and so the search space is huge leading in many cases to overfitting. Deep learning methods have been recently applied to study in a number of issues related to the analysis of gene expression data, including batch correction (Yu et al. 2023), differential expression (Feng et al. 2024), prediction of regulatory networks (Shu et al. 2021) and perturbation prediction (Roohani et al. 2024). However, these also do not utilize prior knowledge leading to weak results in many cases (Song et al. 2024). Another important factor is the small number of patients, especially in phase-2 studies. Many of the methods requires tens of patients to reliably learn patterns and identify differentially expressed genes, but given the scale of phase-2 trials, it can be hard to get required number of patients after subgroup identification.

Graph neural networks (GNNs) (Corso et al. 2024) can be used to integrate prior biological knowledge when learning deep gene expression networks. A gene’s expression in steady-state is a result of its interactions with other genes and GNNs provide the framework to encode such interactions and regulation of biological activity, making them appropriate for modeling biological processes and disease mechanisms. A key advantage of graph networks is their ability to infer the impact of one node in the network on other nodes (for example via regulation or other interactions). This allows for the discovery of nuanced regulatory patterns and context-specific mechanisms that may be missed by other approaches. Their scalability also allows for efficient analysis of large-scale, high-dimensional transcriptomic data without the need for extensive filtering. Indeed, such networks have been used for molecular property prediction (Wu et al. 2023), gene regulatory network prediction (Ji et al. 2024), identifying disease targets (Zhang et al. 2022), clinical risk assessment (Boll et al. 2024), and more. Many flavors of the GNN architectures such as Graph Attention Networks (GATs) and Graph Convolution Networks (GCNs) have been used for various biomedical applications.

In addition to learning context specific deep networks, a number of foundation models (Guo et al. 2025) for gene expression have been introduced recently. Foundation models are trained on large, not related data, and are then fine tuned to process specific datasets (Theodoris et al. 2023). Though foundation models and GNNs have been widely used for different applications, they have rarely been used in the context of modeling timeseries clinical datasets. Combining gene embeddings from foundation models with biological networks through GNNs and applying it to model temporal gene expression can lead to novel insights on genes involved in disease pathology and discovery of dynamic mechanisms associated with disease endotypes.

In this study we leverage gene embeddings from a text based foundation model (GenePT (Chen and Zou 2025)). We finetune GenePT with tissue specific gene expression of healthy samples to capture the context of a specific tissue. We then use the context specific embeddings as part of a GNN to model transcriptomic timeseries datasets. We use GNNs for finetuning the embeddings and for temporal modeling using protein-protein interaction network as the edges in the graph. We applied our method to three clinical time series datasets representing interventions to treat different diseases. As we show, our method accurately recovers known genes/biological mechanisms that underlie disease progression and response and also raises novel hypothesis about mechanisms associated with non-response to treatment. We also show that the proposed method overcomes the important issue of small sample sizes and reliably identifies differentially expressed genes.

## 2 Results

We developed a method for analyzing clinical transcriptomic data using graph neural networks (GNNs). Figure 1 presents an overview of our method focusing on data from ulcerative colitis (UC). To identify genes and pathways that differ between responders and non-responders in UC treatment, we first finetune a gene foundation model (GenePT) (figure 1a) using gene expression of healthy colon data (figure - 1b) by training a GAT to obtain a lower dimensional embedding that captures the context of the tissue (figure - 1d). A temporal GNN architecture is then initialized (node features) with the finetuned embeddings and used to train gene expression of both responders and non-responders. The resulting gene embeddings are then analyzed to obtain insights on genes and biological mechanisms that explain differences in responder and non-responder patients.

**Fig. 1.**
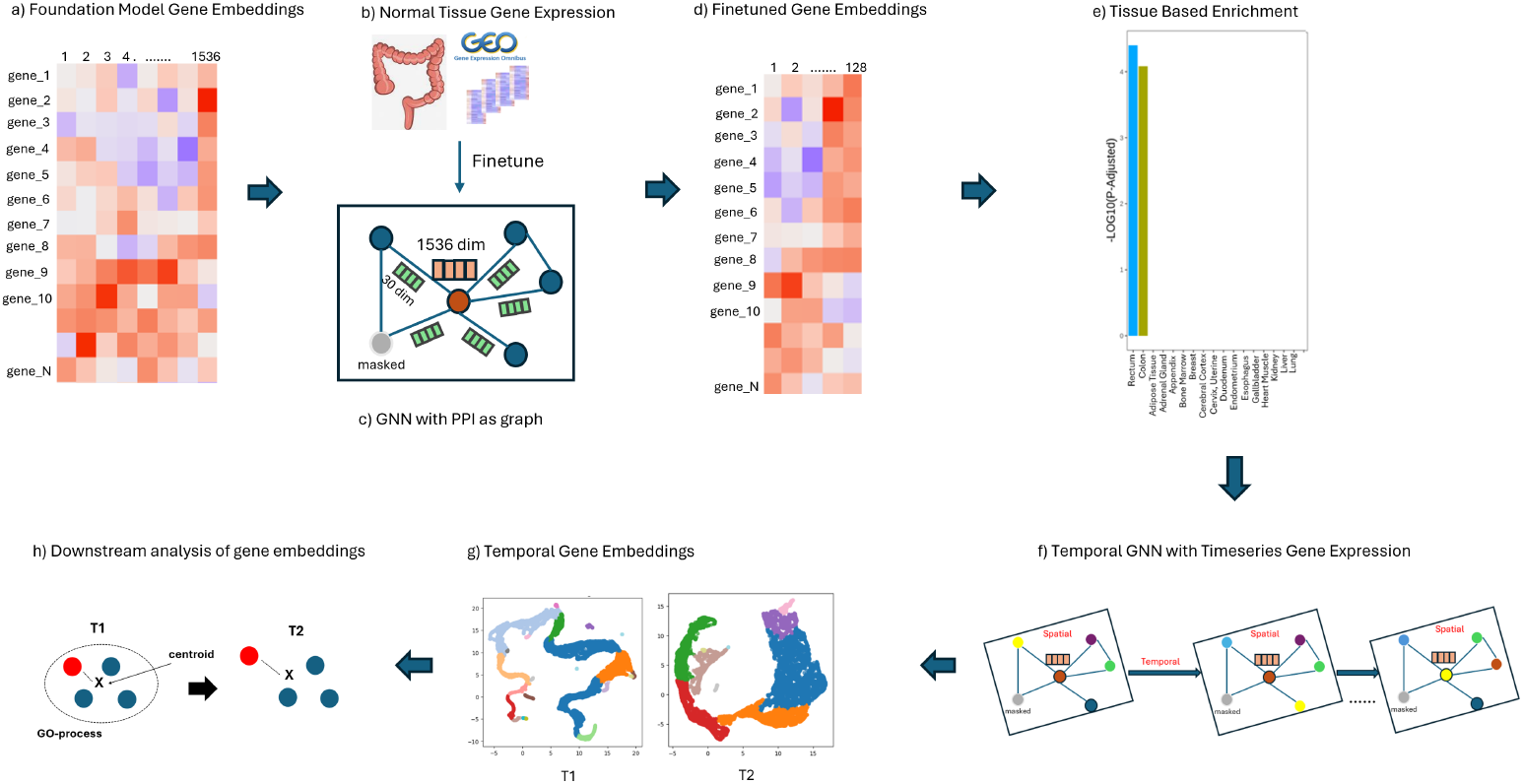
Method overview. a) Gene embeddings from GenePT foundational model with 1536 dimensions and 16000 genes. b) Gene expression of normal samples from the tissue of interest (colon/skin) are extracted from multiple datasets from NCBI-GEO and used for finetuning the GenePT embeddings. c) A GAT is employed to finetune the GenePT embeddings using the PPI as the graph and gene expression from the tissue of interest. d) Finetuned gene embeddings that had a much lower dimensions (128). e) Embeddings finetuned with colon samples are shown to be enriched in rectum and colon tissues. f) The finetuned embeddings are used in a temporal GNN model as node features with TCGN architecture that models gene expression from responder/non-responder datasets. The resulting embeddings are 16 dimensional. g) UMAP representation of the clustered temporal gene embeddings obtained for 2 timepoints shows different neighborhoods for genes. h) Gene embeddings are analyzed by computing the deviation of a gene (red) from the centroid of the GO-Process it is involved in across timepoints T1 and T2.

The method was applied to 3 datasets. Table 1 summarizes the datasets we used and indicates for each the disease, number of responding and non-responding patients, and the timepoints sampled. See Methods for more details.

**Table 1.** Datasets Used in the Study.

| NCBI GEO | Disease | Type<sup>1</sup> | # Patients | Timepoints<sup>2</sup> |
| --- | --- | --- | --- | --- |
| GSE171770 | UC | Resp | 5 | PreTreatment, 4hr, 24hr, 2w, 6w, 14w |
|  | UC | Non-Resp | 4 | PreTreatment, 4hr, 24hr, 2w, 6w |
| GSE171770 | CD | Resp | 4 | PreTreatment, 4hr, 24hr, 2w, 6w, 14w |
|  | CD | Non-Resp | 2 | same as above |
| GSE171012 | Psoriasis | Resp | 9 | PreTreatment, 2w, 4w, 12w |

<sup>1</sup>Resp: Responders or patients who showed signs of remission from the disease following treatment, Non-Resp: Non-Responders or patients who did not show signs of remission following treatment
<sup>2</sup>hr: Hour, w: week

### 2.1 Finetuning gene foundation models

GenePT embeddings were finetuned with the tissue of interest, for example with gene expression from normal colon samples to model timeseries gene expression from ulcerative colitis and Crohn’s patients (Methods). The finetuned embeddings were checked for tissue enrichment and functional enrichment using GO-biological processes. Figure 1e presents the tissue enrichment for colon when using GenePT embeddings after finetuning. As can be seen, the finetuned embeddings are enriched for colon and rectal tissues as expected. The embeddings were further examined for functional relevance, specifically their ability to predict GO-biological processes. To examine the relevance of the embeddings, a logistic regression model was trained on the dimensions of the gene embeddings with GO-Processes as the response variable (Methods). The AUC of the prediction on the test set was 0.64 which is in line with Geneformer embeddings from a recent study (Theodoris et al. 2023). The tissue enrichment of embeddings from the foundation model are distributed across the tissues compared to the finetuned embeddings (supplementary figure 1),

Next we trained separate temporal GNNs for responders and non-responders using the finetuned embeddings as input to the temporal GNN. These embeddings are trained on the gene expression of responders (and non-responders) by masking 1-3% of genes and predicting their expression in the context of the non-masked genes. This enables the network to capture the mechanistic actions of genes that drive the response in the context of their interactions with other genes at each timepoint, where the updated gene embeddings explain the observed gene expression changes (see Figure-4 for details on the architecture). The gene embeddings from the temporal model were used in conjunction with GO-Biological processes to find genes that are variable across timepoints (Methods). Capturing the temporal changes of embedding of a biological process (calculated by averaging embeddings of genes involved) with time provides information on mechanisms that are regulated by the drug used in the study.

### 2.2 Comparing responders and non-responders using temporal modeling

#### 2.2.1 Ulcerative Colitis

Figure 2a presents the biological processes that were identified for UC responders and non-responders. For responders we see that many of the mechanisms that are known to be associated with wound healing are correctly identified as up regulated. These include aerobic electron transport chain (GO:0019646), retinol metabolic process (GO:0042572), lung epithelium development (GO:0060428), action potential (GO:0001508), retinoic acid metabolic process. These pathways are reflective of the mucosal healing of the gut following response to treatment, an important aspect the disease modification in IBD. In responder patients, the interleukin-6-mediated signaling pathway (GO:0070102) is down regulated confirming anti-IL6 blockade. Additional immune response mechanisms such as interleukin-15 signaling pathway and production of IL-1b, IL-2 is down regulated in responders. This is potentially reflective of blockade of the feed forward loop of IL-6/ IL-1b/ IL-15 and JAK-STAT pathway primarily modulated by STAT3 (Mori et al. 2011), and reduced IL-2 signaling, potentially reflective of return of the T cell activation and Treg function to steady state. In contrast, in non-responder patients, although interleukin-6-mediated biology is down regulated, some potential immune related mechanisms of resistance to therapy like B cell receptor signaling pathway (GO:0050853) are elevated. Notably, healing mechanisms such as wound healing, spreading of cells (GO:0044319), epithelial cell-cell adhesion (GO:0090136), calcium-dependent cell-cell adhesion via plasma membrane cell adhesion molecules (GO:0016339) are down regulated in non-responders.

**Fig. 2.**
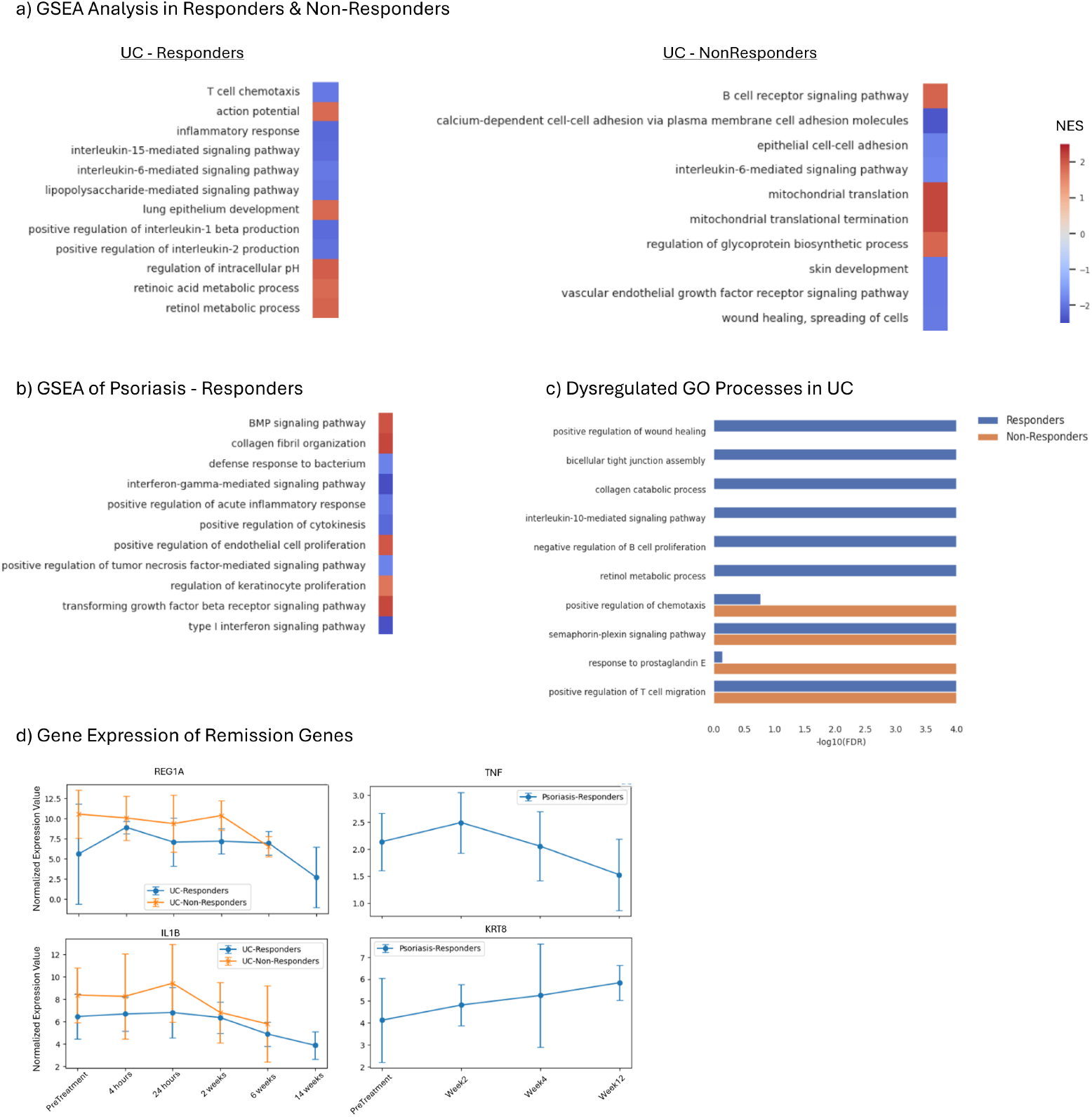
Comparison of Responder and Non-Responders. a) GSEA results of responder and nonresponder groups in ulcerative colitis. A subset of significant GO-Biological processes are shown with the Normalized Enrichment Score (NES). b) GSEA results of responder group for Psoriasis. A subset of significant GO-Biological processes are shown with the Normalized Enrichment Score (NES). c) GO-Biological processes obtained form the perturbation analysis in responder and non-responder groups in UC are shown. The -log10(FDR) values are plotted for the 2 groups. FDR *<* 1e-4 were truncated to 1e-4 for display purposes. d) Boxplots of gene expression for remission-related genes that are detected by the method are shown for UC and Psoriasis. The bar represent the standard deviation across samples at a given timepoint.

Results of GO-embedding perturbation (see methods) analysis (2c) shows mechanisms that are changed with time in responders and non-responders. Although, these do not have directional information, it is important to understand mechanisms that change with time. Mechanisms related to remission such as positive regulation of wound healing, bicellular tight junction assembly, collagen catabolic process, interleukin-10-mediated signaling pathway, and negative regulation of B cell proliferation are dysregulated only in responders. In contrast, positive regulation of chemotaxis, and response to prostaglandin E are dysregulated exclusively in nonresponders. Semaphorin-plexin signaling pathway that is associated with dendritic and T-cell activation both change in both responders and non-responders, although directionality is uncertain.

#### 2.2.2 Crohn’s Disease

Figure-2 in the supplementary file lists the results from GSEA analysis in responders and non-responder patients. Mechanisms in responders such as endodermal cell differentiation (GO:0035987) and positive regulation of lipid kinase activity (GO:0090218) are up regulated that point to healing activity, while negative regulation of interleukin-6 production (GO:0032715) is also up regulated that points to decreasing activity of IL-6 signaling confirming drug’s mechanism of action. Interestingly mitochondrial activity - mitochondrial gene expression (GO:0140053) and mitochondrial translational elongation (GO:0070125) is down regulated pointing to decreased metabolic activity which may be a result of mucosal healing and return to homeostatic immune status. In non-responders, immune related activity linked to positive regulation of interleukin-2 production (GO:0032743), cellular response to interferon-gamma (GO:0071346) potentially implicate a disrupted barrier with ensuing inflammation driven by microbial super-antigens and type 1 immune response; chemokine-mediated signaling pathway (GO:0070098) resulting from increased inflammatory response leading to a positive fee-forward loop of increased inflammation and tissue damage and positive regulation of interleukin-2 production (GO:0032743) as a potential mechanism of increased T cell driven pathogenesis as well as increase Treg response as an attempt to reduce the inflammation. Mechanisms related to protein synthesis activity are down regulated, potentially reflecting dysfunctional repair mechanisms in non responders.

#### 2.2.3 Psoriasis

GSEA results for Psoriasis (Figure 2b) that consisted of only responders 2a due to the know depth of efficacy of Secukinumab in this disease reaching PASI-75 and PASI-90 in high proportion of patients (Yang et al. 2018). Notably, BMP signaling (GO:0030509), TGFb signaling (GO:0007179), positive regulation of endothelial cell proliferation (GO:0001938) and regulation of keratinocyte proliferation (GO:0010837) (Li et al. 2024) that are associated with resolution of inflammation and skin healing are up regulated. GO processes related to cellular proliferation and immune response related mechanisms for innate and adaptive such as defense response to bacterium (GO:0042742), interferon-gamma-mediated signaling pathway (GO:0060333), positive regulation of tumor necrosis factor-mediated signaling pathway (GO:1903265) and others.

Figure 2d shows plots of gene expression of 4 genes of interest in UC and psoriasis that are not captured using differential expression analysis in the original publications of these studies or by prior methods. One of the main reasons these genes were not detected previously is because of the variance induced by the heterogeneous patient population and subtle temporal changes that are hard to pick through traditional methods. The ability of our method to utilize embeddings that smooth such variability while still maintaining overall response patterns allows it to correctly identify these genes.

### 2.3 Differential networks between responders and non-responders in UC

We focused on gene embeddings that showed a significant change following treatment with Olamkicept in responders but not in non-responders. To extract the sub-network for a gene, we extracted the 1-hop (immediate neighbor) genes and connections among them from the PPI network. If the sub-graph was too large (*>* 200 genes), then we restricted it to genes that are differentially expressed (pval *<* 0.05) in either responders or non-responders so that the the figure is interpretable. The directionality of regulation was added to the genes, which was calculated as the average log-fold change of genes from the samples of last timepoint compared to pretreatment samples.

Figure 3a shows that CD19 network is significantly impacted in responders with reduced CD38 expression and B cell receptor signaling components like VAV1, FYN, LYN, and SYK implying reduced B cell activation and reduced plasmablast/plasma cell phenotype. CXCL13 is also reduced potentially reflecting reduced B cell and TFH migration and reduction in tertiary lymphoid organs (Hui et al. 2024). B cells expansion and activated B cell states have been recently shown to impair epithelial-stromal cell interactions required for mucosal healing in Ulcerative colitis (Frede et al. 2022). Interestingly, some of the genes like CD19, MS4A1, CXCR4, SCARB1, SCARB2 show elevated expression which might reflect predominance of the naïve B cell compartment in responders and restoration of the homeostatic mechanisms of the gut epithelium related to mucosal healing. Also the significant chemokine network change in responders vs non-responders reflects a less inflammatory state of the epithelial and stromal cell compartments leading to reduced production of pro-inflammatory chemokines thereby reducing the inflammatory load of the gut (Singh et al. 2016).

**Fig. 3.**
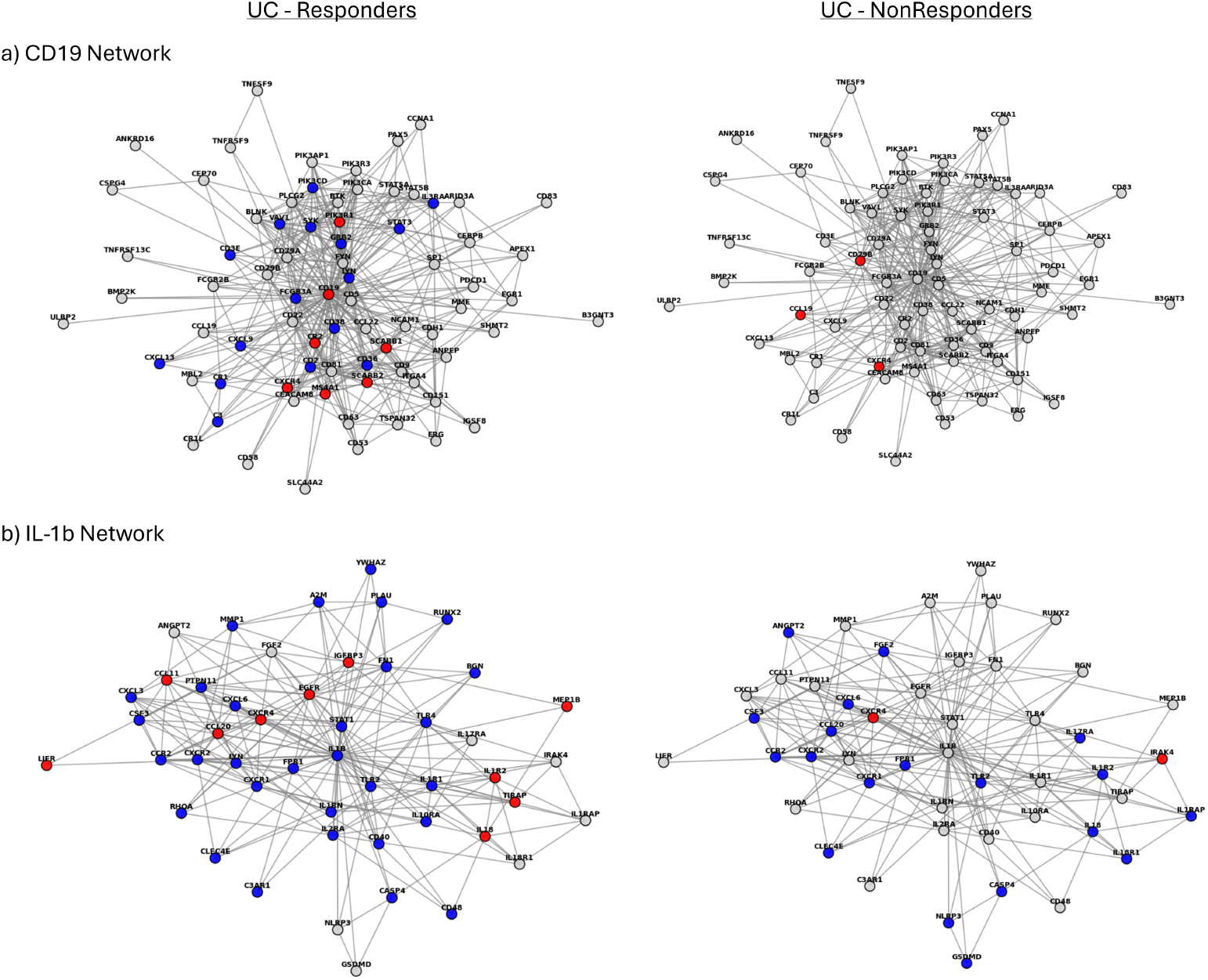
Network Neighborhood of IL1b, CD19 in UC responder and non-responder groups. The up regulated (log-fold change *>* 0) genes are shown in red while down regulated (log-fold change *<* 0) genes are shown in blue. a) 1-hop neighborhood of CD19 (center of the figure 3a). b) 1-hop neighborhood of IL1b (center). Only the differentially expressed genes in either responder and nonresponder groups are shown.

Next, we explored the neighborhood of IL-1b Figure 3b which is one of the proinflammatory factors in UC and CD (Aggeletopoulou et al. 2024) and contrasted it between responder and non-responder groups. It consisted of *>* 200 genes many of which were not differentially expressed, consequently only genes that were differentially expressed (p-value *<* 0.05) in either responders or non-responders are shown. It can be seen that in the responder group the IL-1b (center gene) is down regulated and its neighborhood which includes chemokine receptors such as CXCR1/2, genes related to T-cell activity (CD40, CD48) along with STAT1 are mostly down regulated. In contrast, in the non-responder group IL-1b is not down regulated and only a few of its neighbors are down regulated.

### 2.4 Comparison with Previous Approaches

Table 2 presents comparison of the proposed approach (Proposed Method) with the state-of-art time series methods such as MASigPro, TiSA and a recently developed constrained pseudotime ordering method (Pseudo-Ord) for responders in UC, CD and Psoriasis. The comparison is made on the ability of the methods to detect differential expression (DE) at FDR *<* 0.05 and detect primary treatment mechanisms. The proposed method consistently detects higher number of differentially expressed genes compared to other methods. MASigPro and TiSA detect many genes with p-value *<* 0.05, but very few after adjusting for multiple hypothesis correction. Pseudo-ordering method detects very large number of genes in UC and psoriasis, but fails to detect any in CD because of low sample size.

**Table 2.**
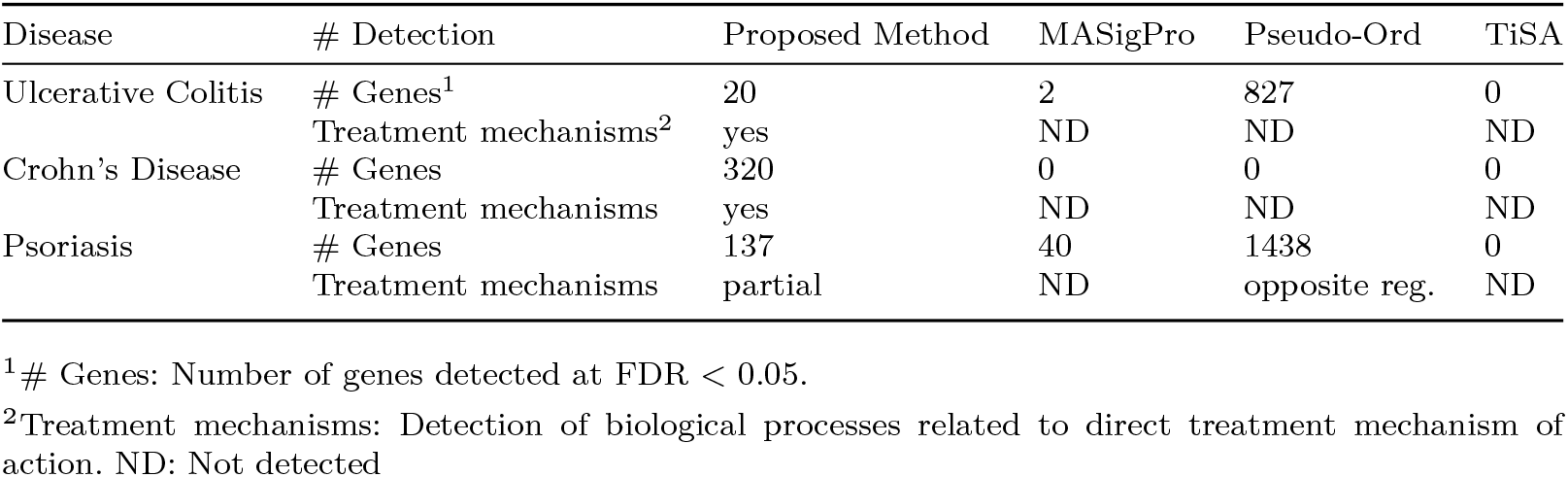
Comparison with Previous Methods with Responder Patient Group.

The proposed method also detects the treatment mechanism of action for UC and CD. For example in IL-6 treatment for UC and CD, one of the down regulated pathways is interleukin-6-mediated signaling pathway (GO:0070102) which is directly related to the mechanism of action of olamkicept that blocks IL-6 trans-signaling. Healing mechanisms such as retinoic acid metabolic process (GO:0042573) which is activated as a result of the drug are also detected in UC responder subgroup. For psoriasis, patients were treated with secukinumab that disrupts IL-17 pathway and should be detected as down regulated. Pseudo-ordering method detects positive regulation of interleukin-17 production (GO:0032740) with a positive normalized enrichment score which is contrary to the expected outcome. The proposed method is unable to detect IL-17 signaling but detects down regulation of *interferon-gamma-mediated signaling pathway (GO:0060333)*, and *positive regulation of interleukin-23 production (GO:0032747)*.

As for treatment related genes, our method correctly identifies differential expression of such genes in all 3 studies. Figure 2d shows remission-related genes for UC and psoriasis in responder and non-responder subgroups. The normalized genes expression is shown along with the standard deviation at each timepoint, which is quite large owing to heterogeneity in human samples. In contrast, MASigPro and TiSA fail to detect significant genes in these studies due to large standard deviations within different timepoints. For example, even though the mean expression values of REG1A, IL-1b in UC responder and non-responder subgroups are different across timepoints, their standard deviations overlap makes their detection difficult for prior methods. Interestingly, TNF has a small changes in expression across time, but plays a pivotal role in reduction of inflammation was detected by the proposed method.

### 2.5 Ablation Studies

We conducted ablation studies to ascertain the methodology we used in this paper. For this, we tested the method at 3 critical junctures, 1) use of PPI within GNNs to model gene expression 2) finetuning of foundation gene embeddings, and 3) use of embeddings for differential expression as opposed to gene expression.

#### 2.5.1 Use of PPI Network

We first performed analysis to test if the use of prior biological data as prior to the GNN helps in modeling gene expression. To check if the PPI information improves the learned neural network we compared the performance of the model using the actual PPI and a model that uses randomized PPI with the same number of edges (Methods). To quantify the difference in the ability of the network to understand gene expression we masked a subset of genes for both (actual and randomized) networks and used the corresponding model to predict gene expression. Table 2 shows the Rsquared obtained on the test set across the 3 datasets for both settings. As can be seen, model fit as measured by R-squared when using the real PPI is higher compared to the models created with perturbed PPI across all datasets (0.72 vs 0.16 for UC). This shows that the PPI has meaningful information and provides biologically relevant priors for modeling gene expression.

#### 2.5.2 Finetuning of Embeddings

The foundation embeddings were finetuned using gene expression from the tissue of interest. To check relevance of finetuning, we compared our results to results obtained by using embeddings from GenePT directly. Again, we masked genes in the time series data and used the model to predict them. As can be seen, the R-squared of the finetuned model are consistently higher across the 3 studies when compared to the original GenePT embeddings, 0.72 vs 0.51 for UC, 0.58 vs 0.18 for CD, and 0.53 vs 0.23 for psoriasis. This demonstrates that finetuning plays an important role in modeling temporal gene expression data.

The tissue enrichment of the GenePT embeddings and that of the skin can be seen in supplementary Figure-1a,b. GenePT embeddings as expected are enriched in many different tissues and in contrast the finetuned embeddings obtained by training with healthy skin samples are enriched in skin and adipose tissues.

#### 2.5.3 Embeddings vs. DE analysis

Embeddings from the temporal GNN model were used to find functionally important genes by analyzing the deviation of gene embeddings over time (see methods). To test if such a strategy improves on standard DE analysis we compared it to a method that only uses differential expression to identify significant genes. Results presented in Table 2 show that DE analysis does not identify significant genes in most datasets likely because of noisy and small datasets. We also tested our strategy of using deviations of embeddings as described in the methods section, and used gene expression in lieu of embeddings. This also did not detect any genes related to treatment mechanism across the three datasets.

## 3 Discussion

Clinical transcriptomic data offer tremendous opportunity for drug discovery through back-translation (Shakhnovich 2018) but pose a unique challenge of increased heterogeneity in gene expression because of varied demographics, concomitant medications taken by patients, genetic variation, epigenetic factors and more. Another problem is the size of datasets. Although there are some large observational studies, clinical studies including phase-2 studies profile only a small number of individuals. Understanding the differences between patients that respond to the drug (responders) and those that do not (non-responders) is critical for determining the mechanisms driving disease in heterogeneous populations. Recent methods relied on concepts from single cell trajectory analysis such as constrained pseudo-ordering (Mathur et al. 2024; Campbell and Yau 2018) to remove the unwanted variation by placing samples on a disease/treatment axis. However these methods do not incorporate prior knowledge such as gene-gene regulatory interactions, which has been shown to be very useful in modeling gene expression (Ji et al. 2024).

In this paper, we have developed a method for analyzing time series data from clinical transcriptomic studies that extends gene-based foundational models, utilizes normal tissue expression from NCBI-GEO, and relies on prior information using PPI networks. We first finetune gene embeddings from an LLM (GenePT) on relevant healthy tissue data figure 1. Next we use the gene embeddings from the finetuned models together with the gene expression from the clinical studies to learn a time series Graph Neural Network where edges represent gene-gene interactions. Our method utilizes BERT-style training (Devlin et al. 2019) leading to impact from both the local neighborhood and the entire graph on the learned embeddings. Gene/node embeddings are updated using the embeddings of their immediate neighbors (local neighborhood) and also based on the larger network. This ensures the gene embeddings are representative of the gene expression in the context of the connected components in the PPI network. Another important aspect is to finetune the foundation embeddings. Applying our method to responder and non responder data from several clinical trials shows that it is able to identify key genes and pathways that drive response and that our method provides novel hypotheses for mechanisms affecting treatment efficacy.

GNNs work through information flow and use the local and global neighborhood to update the node information. This mimics the information flow in biological networks and hence serves as promising architecture to model biological mechanisms. Temporal GNNs have been used to model time series data for other domains but have not been widely used for time series expression data (Giovanoudi and Rafailidis 2025). Temporal gene embeddings (embeddings at each timepoint) represent the gene context in response to perturbation (treatment) at each timepoint. These are rich in information, but lack directional context and magnitude of change. In fact much of the previous work involved using embeddings to predict connectivity between genes or for classification purposes. To overcome these limitations, we used the deviation of a gene between successive timepoints in the time series and used a null model to assess if the deviations were significant by computing a pvalue. For directionality, we used the fold change between the baseline and the timepoint at which the clinical remission assessment was made. A composite score of gene significance and direction of regulation was fed into GSEA to obtain mechanisms involved in the clinical response study. Given gene embeddings, a GO-Process centric view was also created by averaging the embeddings of genes belonging to the process.

A number of genes that are identified by our method for UC responders (full list in supplementary file) are known to be linked to treatment response. For example CCL20 attracts Th17 cells to the inflamed area, while CXCL9, CXCL10 and CXCL13 recruit T and B cells. REG1A, REG1B, REG3A are involved in epithelial regeneration. Many of these aforementioned genes along with ZAP70, IL-1b, CD2, CD19 have been associated with UC. Additionally a few interesting genes such as CA3 (Okada and Ikemoto 2022) have been recently shown to be a biomarker for UC as an immune regulator. THSD7B and FAM151A are novel predictions that have not been found to be associated with UC in the current literature. In Crohn’s disease, our method detected several genes that have been previously associated with the disease (full list in supplementary file). These include T-cell activation genes such as CD2, CD247, ZAP70; matrix metalloproteinases; involved in tissue remodeling and barrier breakdown in CD, and COL1A1, COL3A1, COL5A1, COL6A3 that are involved in extracellular matrix generation. Other interesting genes are ADM5 which is known to be an antiinflammatory peptide (Ashizuka et al. 2021), GALNT15 which is an O-glycosylation enzyme that is linked to mucous production and FAM151A which was also found in UC responders among many other genes.

Much of the previous work in timeseries expression analysis focused on fitting different models for gene expression between the first and last time points or on finding differential expression between successive timepoints. Indeed, in the original studies for all the data presented in this paper findings were based on comparison of samples at pretreatment and the last time point or between pre-treatment and all time points. Given the heterogeneity of clinical transcriptomic data, a method that transforms gene expression that reduces unwanted variation is much better suited to capture subtle differences. Results (Figure 2, Table 2) indicate that our method seems to achieve such regularization. Using gene expression directly, as is done in MASigPro and TiSA, fails to detect genes at reliable thresholds (FDR *<* 0.05) and does not lead to the detection of treatment-related genes. Constrained pseudo-ordering that uses fitted gene expression detects genes in UC and psoriasis. The GNN architecture lays more emphasis on message passing between connected genes, and in the process removes unwanted variation. Consider the Crohnś disease dataset that has very few samples (4) per timepoint in the responder group. All prior methods failed to identify relevant biological mechanisms or detect differential expression. In contrast, by constructing context specific finetuned models over time our method was able to overcome this key limitation. Figure 3 shows that multiple genes change in the immediate neighborhood of 2 key genes that are associated with pathology of UC. The transfer of information to a gene from its neighborhood enables the gene to capture the context of gene expression across its neighbors along time and this helps overcome the heterogeneity associated with gene expression.

Determining the dimensionality of embeddings is not an exact science, but is mostly based on experimentation (Purchase et al. 2022). We experimented with different dimensions of embeddings (8, 16, 32) and used PCA to check if there are too many redundant dimensions. For the finetuned embeddings the original Gene-PT embeddings were 1536 dimensions and we reduced it to 756, 256, and 128 dimensions. We used tissue enrichment as a criterion to decide on the length of the embeddings (more info in the supplementary file). For the temporal embeddings we used PCA and settled on the length of 16 (supplementary figure-1c,d). Ablation studies were used build confidence into the methodology at crucial stages. Results from Table 3 show that inclusion of PPI network as a graph is beneficial to modeling gene expression and has the highest impact on performance of the model. Finetuning is also shown to be very important especially when the number of samples is low in the case of Crohnś disease. Although the proposed method offers advantages over existing frameworks, it heavily relies on the underlying graph structure for message passing. Human gene annotation is biased (Haynes et al. 2018) towards well studied genes and this can limit the discovery of novel genes associated with disease. Genes that are not part of the graph structure or in disconnected components are left out of the analysis. Future work should attempt to overcome the limitations of disconnected graphs by using additional interaction sources and extend the framework/architecture to include clinical scores and incorporate pseudo-ordering to overcome the issue of disparate disease states of patients during a visit.

**Table 3.** R-Squared of Models from Ablation Studies on Test Sets.

| Disease | Dataset | Actual<sup>1</sup> | PPI<sup>2</sup> | Embeddings<sup>3</sup> |
| --- | --- | --- | --- | --- |
| Ulcerative Colitis | Responders | 0.72 | 0.16 | 0.51 |
|  | Non-Responders | 0.66 | 0.23 | 0.28 |
| Crohn’s Disease | Responders | 0.58 | 0.14 | 0.18 |
|  | Non-Responders | 0.57 | 0.23 | 0.44 |
| Psoriasis | Responders | 0.53 | -0.04 | 0.23 |

<sup>1</sup>Actual values from the Method.
<sup>2</sup>Randomized PPI network used to test use of PPI.
<sup>3</sup>Testing the foundation embeddings instead of tissue-finetuned embeddings.

## 4 Conclusion

We have developed a novel method to model clinical timeseries expression data that overcomes excessive gene variation and small sample size issues in transcriptomic clinical datasets. It leverages foundational models, finetuning, and prior interaction networks to uncover known and novel mechanisms associated with response and nonresponse to drug treatment. Some of the limitations are biases or lack of annotation in the graph structure and lack of directionality of gene embeddings. Future work includes extending the framework to include clinical information and use of pseudo ordering of patients rather than relying on visit information alone.

## 5 Materials and Methods

### 5.1 Datasets and Preprocessing

We collected 3 disease-related datasets for testing our method. The first two, Crohn’s and ulcerative colitis (UC) datasets were obtained from GSE171770. This dataset is from a Phase 2a trial for the drug Olamkicept (Schreiber et al. 2021) on IBD (Inflammatory Bowel Disease) patients, 7 patients each for UC and CD, in which the patients received the drug for 14 weeks and samples from the intestinal mucosa were collected at pretreatment (0hr), 4 hours, 24 hours, 2 weeks, 6 weeks, and 14 weeks. Clinical scores (Mayo score for UC and CDAI - Clinical Disease Activity Index for CD) were used to determine responder and non-responder patients at 14 weeks. Gene expression data for the UC Non-Responder population was not available in the NCBI-GEO dataset. The third dataset is a Psoriasis (GSE171012) dataset in which 15 patients were treated with secukinumab, an IL-17 inhibitor and samples were collected at pre-treatment, 2 weeks, 4 weeks, and 12 weeks. The PASI (Psoriasis Area Severity Index) clinical scores were used to determine that all patients had a progressive reduction in severity of the disease and all patients were deemed as responders. Patients that had samples from all timepoints were selected. Multiple normal tissue datasets for colon and skin were downloaded from NCBI GEO. The list of datasets are shown in the supplementary Table-1.

Raw gene counts were obtained from the supplementary files in NCBI GEO for the three RNASeq datasets (Psoriasis, UC, CD) and normal tissues. Only protein-coding genes that had at least 5 counts in at least 1% of the samples were kept. In the case of duplicated gene identifiers, the gene with highest mean expression was considered. The datasets were then normalized using TMM (Robinson and Oshlack 2010). This was used as an input to ComBat (Johnson et al. 2007) batch correction to obtain batch corrected expression values.

Protein-protein interactions (PPI) data was obtained from StringDB and was supplemented with protein-DNA interactions (Transcription Factor-Targets) (Zhang et al. 2020) to account for gene regulatory interactions. The PPI was filtered using genes that were expressed in the tissue of interest and the maximal connected graph was extracted to be used in the GNN.

### 5.2 GNN Architectures

We used two GNN architectures, one for finetuning the gene embeddings from the foundation model and the second as a basis for our temporal architecture to model gene expression from the responder and non-responders over time. Separate temporal models were created for the responder and non-responder groups. Nodes in these models represent genes in the protein-protein interaction network.

The finetuning GNN was based on Graph Attention Networks (GATs) (Boll et al. 2024) architecture that allows the model to implicitly learn the importance of different neighbors for a given node, rather than relying on predefined static weights. The edge attributes consisted of concatenated vectors of gene expression of the 2 genes that shared the edge. The GAT model consisted of 3 layers with decreasing sizes (756, 256, 128) of node embeddings (figure 4a), transforming the 1536 dimensional foundation gene embeddings to 128-d embeddings. RELU was used as the non-linear function throughout the model and consisted of a single linear layer before prediction. GAT consisted of a single attention head. Gene expression from up to 15 normal samples from 180 samples (in colon) were used in each iteration during training. The training was done by masking 1-3% of the genes (nodes) by setting their embeddings to 0, and predicting the gene expression for all genes. The training error corresponding to the masked genes was back-propagated. Different set of genes were masked at each epoch and the model was trained for 200 epochs.

**Fig. 4.**
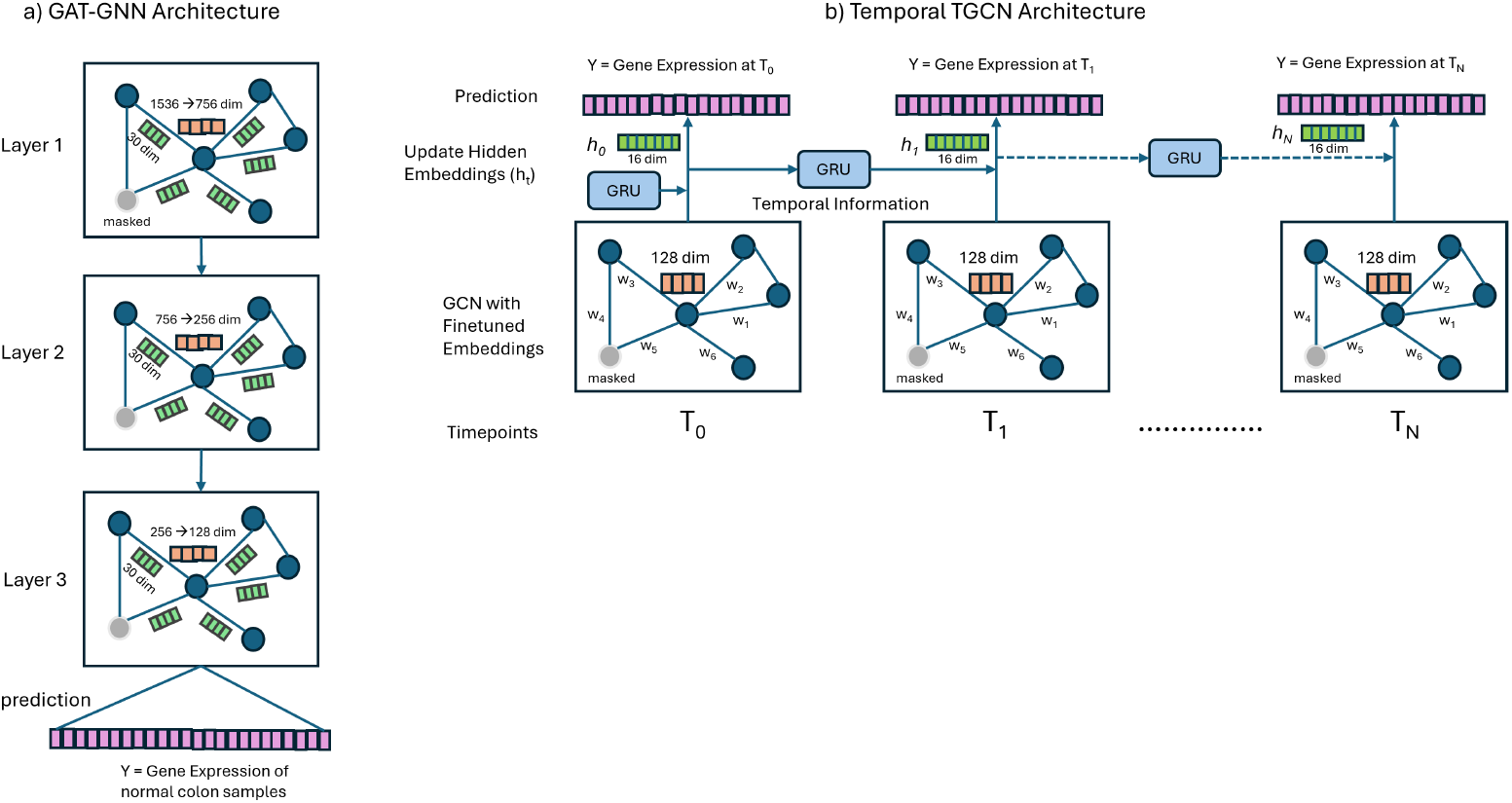
GNN Architecture for Finetuning and Temporal Modeling. a) Shows a 3-layered GNN-GAT architecture with 1536 dimensional node embeddings and a 30 dimensional edge embedding. The node embedding dimensions are reduced with the number of layers. A set of nodes (genes) are masked for training. b) Illustrates the temporal TCGN architecture with GCN capturing the spatial relationships and Gated-Recurrent Unit (GRU) capturing the temporal information. The network is initialized with the 128 dimensional finetuned embedding. Same set of nodes are masked at all timepoints. The resultant embedding is the transformed 16-dimensional embeddings (hidden state) obtained from GRU and GCN.

Given graph *G*, embeddings *E* ∈ *R*^*n×d*^, where *n* are genes, *d* dimension of the embedding, the GAT model predicts the estimated gene expression values *ŷ*.

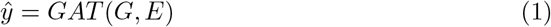

1% of the genes were selected for the test set and were not used during training. The R-squared value was calculated on the test set. The finetuned 128-d embeddings were extracted from the nodes after the training. We experimented by masking between 1-20% genes and decided to keep 1-3% based on the training error as higher percentages of masked genes gave a large training error and appeared to be significantly underfitting.

These finetuned embeddings should capture the context of gene expression of the tissue. To determine if they indeed capture tissue relevant biology we projected these embeddings into 2-d space using UMAP and clustered them. These clusters were checked for tissue enrichment using TissueEnrich (Jain and Tuteja 2018) and as we discuss in Results the finetuned embeddings were enriched for the relevant tissues - Figure 2e.

Next we used the finetuned 128-d embedding as the node embeddings/features for the temporal GNN response model. The TGCN (Zhao et al. 2019) architecture was used to model the timeseries gene expression of responder and non-responder patients. Separate models were created for each group. TGCN consists of a series of graph convolutional networks (GCN) that are connected with a gated recurrent unit (GRU) to carry the temporal information across the layers. Figure 4b shows the details of our architecture in which the finetuned 128-d embeddings are used as node embeddings. The temporal information is merged with the graph convolution output to compute the hidden state, which is then connected to a fully connected layer for predictions. The edge weights between a pair of genes at a given timepoint were a function of the distance between the gene expression values. The distance between a pair of genes is calculated using the euclidean distance, as shown below.

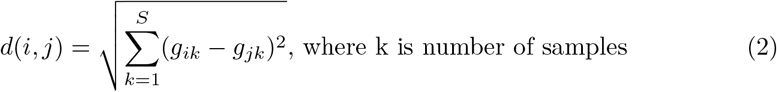

Since the edge weights represent similarity, the distances were converted into similarity scores by normalizing the pairwise distances with the maximum euclidean distance, *d*_*max*_, which is the maximum distance between any pair of genes. The edge weight between a pair of genes *i* and *j* is calculated by

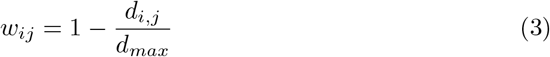

The finetuned embeddings matrix *E* ∈ *R*^*n×m*^, where *n* is the number of genes and *m* is the dimension of the embedding, was constructed as follows:

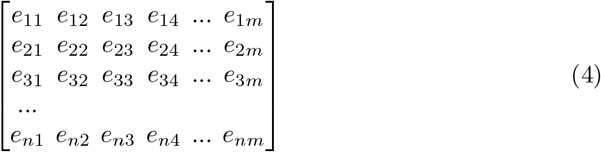

where *e*_*ij*_ represents *j th* dimension of gene *i* in the finetuned embedding. The gene expression matrix *G*^*t*^ at time *t* is given below, where *n* are the number of genes and *k* is the number of samples

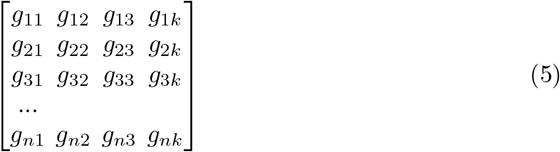

To account for samples from different patients at a given timepoint, the finetuned embeddings were scaled and averaged with the gene expression of samples at each timepoint resulting in *E′* given by

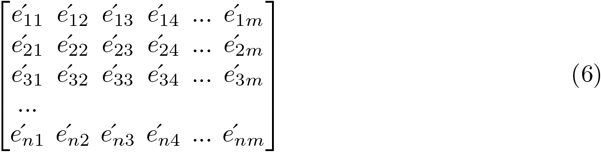

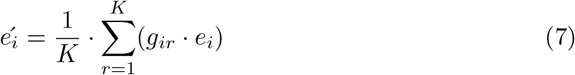

where K is number of samples at timepoint *t, e*_*i*_ = [*e*_11_, *e*_12_, …*e*_1*m*_], and 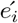 is the scaled-averaged embeddings across *K* samples at time *t*. These finetuned and scaled embeddings were used in the temporal model as node (gene) embeddings. Note that the finetuned embeddings were scaled with gene expression according to the timepoint resulting in different scaled embeddings at each timepoint.

Scaling and averaging the embeddings across *K* samples ensures that the resulting node embedding for each gene at a given time point reflects both the foundational knowledge from the pretrained embeddings and the actual gene expression patterns observed across all patients. This approach helps reduce the impact of noise or outliers from individual samples, yielding a robust and representative embedding for downstream temporal modeling.

The training was carried out by masking 1-3% genes and predicting the gene expression of samples at a given timepoint *t*. For example, if the dataset had 4 timepoints, then the model was trained by predicting the gene expression values for the patient samples at each *t*. The genes were masked for all timepoints and their expression values were predicted for all patients. The root mean-squared error was calculated on the masked genes and was used in backpropagation. The information learned at timepoint *t* is carried over to the next timepoint (*t* + 1) using the GRU, where the same set of genes are masked again. The hidden state is computed using the information from GRU and output from the graph convolution. The hidden state is then fed into a linear layer with RELU activation function to predict gene expression at (*t* + 1). These hidden states at each timepoint constituted the embeddings of the model and represent the temporal embeddings. Temporal embedding at time *t* is given by *H*^*t*^.

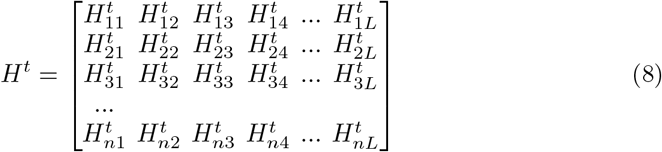

### 5.3 Significant Gene and GO-Process Identification

Temporal embeddings were used to find genes that changed over time by comparing the embeddings at subsequent timepoints. Given the initial timepoint *t*, the deviation for a gene was calculated as the euclidean distance between the gene’s embeddings at time *t* + 1 and *t*, represented as 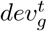. The subsequent deviations were calculated between embeddings between times *t* + 1 and *t* + 2. The total deviation *dev*_*g*_ was the sum of all deviations for the gene *g* as shown below.

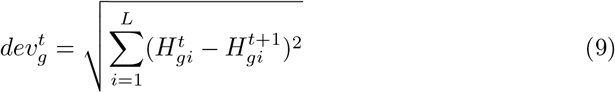

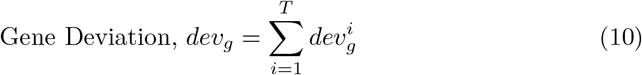

To find if the total deviation for a gene is significant, we generated a null distribution of the temporal model. This is achieved by randomizing the edge connections in the PPI network and training the neural network with gene expression the same way as for the responder and non-responder models. Specifically, the model is trained on randomized PPI and embeddings are constructed while regressing on the actual gene expression values. This was done 1000 times to generate a null distribution of temporal embeddings separately for responder and non-responder patients.

The p-value for a gene is calculated by comparing its total deviation to the null distribution as shown in Equation 10. It is the fraction of random deviations of gene *g* greater than the actual deviation.

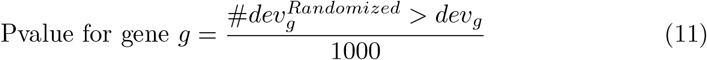

Since gene embeddings lack directionality, the gene expression values were used to determine the direction of regulation. Specifically, the log-fold change was calculated between the last timepoint and the first (pretreatment), with the later serving as the baseline. The gene was deemed to be up regulated it it had a positive fold change and down regulated otherwise. Thereupon, the rank of genes was calculated for GSEA (Subramanian et al. 2005) using, *rank*_*g*_ = *Regulation*_*g*_ *log*_10_(*pvalue*_*g*_). GSEA was conducted using GSEApy (Fang et al. 2023).

We also compute the GO-Process deviation to check if the GO-Process deviates with time. The GO-Process deviation was computed as the average deviation of gene embeddings that are associated with the GO-Process, also referred to as the centroid of the genes in a GO-Process. Thereupon, the deviation of a GO-process was calculated the same way as gene deviation by finding the euclidean distance between centroid of GO-Process embeddings at subsequent timepoints and aggregative their deviations. The null distribution was computed the same way as for genes, but with GO-Processes and subsequently p-values were calculated. The adjusted p-values for both genes and GO-processes were calculated using FDR. It is important to note that the identified GO-Processes lack directionality. It should be noted that for genes, the p-value has tight scale given only 1000 simulations were used in the null distribution and the smallest p-value to resolve is 1/1000. This compression of p-values may lead to FDR being less precise for low p-values.

### 5.4 Procedure for Ablation Studies

Ablations studies were conducted for assessing the relevance of different parts of the methodology - use of PPI network, finetuning of foundational embeddings, and using the deviation of gene embeddings. To check if the PPI network improves gene embeddings when modeling gene expression, we randomized the PPI network such that the degree of connectivity of each node was preserved. The randomized PPI network was then used in the GNN along with the gene embeddings to model the gene expression data by training the model to predict gene expression values and masking 1-3% genes. R-squared was recorded for the test set and repeated 10 times to measure stochasticity of the model fit. These randomized networks also serve as the baseline for performance. To assess if it is necessary to finetune the foundational embeddings before using them in the temporal model, we used the foundational model embeddings directly for training the temporal model and compared the R-squared of the test set with that of the model that used finetuned embeddings. Finally, to assess the utility of using of gene embeddings instead of gene expression, we used the methods described to compute gene deviation using gene expression and compared the findings with those of temporal embeddings.

### 5.5 Implementation of Previous Methods

The method was compared with several prior methods (TiSA, MASigPro, constrained pseudo-ordering) on all 3 datasets. MASigPro and pseudo-ordering methods require a control, so in case of UC, CD, and psoriasis the untreated patient samples were used as controls. MASigPro requires at least 3 replicate samples and hence could not be run on the CD dataset. TiSA makes pairwise comparisons at subsequent timepoints, so there would be 5 comparisons for 6 timepoints in UC and CD. Genes that were detected to be differentially expressed in at least 3 pairwise comparisons were selected. The R-packages for TiSA and MASigPro were used with the default parameters to obtain genes that were differentially expressed (FDR *<* 0.05). Python package for constrained pseudo-ordering was used to calculate the pseudo-ordering of samples and clustering was applied to obtain pseudo-timepoints. As indicated in the paper, singletons were discarded (Mathur et al. 2024) and the clustered timepoints were used as pseudo disease states. Even though the method incorporates clinical scores, only the gene expression values were used to make a fair comparison across all 4 methods. Since the pseudo-ordering method lacks a framework to calculate differential expression across time, differential expression was calculated at each pseudo-timepoint by comparing the fitted gene expression values with that of the pretreatment samples as was done in the paper. Genes that were differentially expressed at all pseudo-timepoints were extracted. GSEA analysis was done at the last pseudo timepoint to obtain enriched GO-processes (FDR *<* 0.05).

### 5.6 Use of LLMs

LLMs such as chatGPT-4o, Gemini-2.5, and Perplexity were used to understand pathways related to the mechanism of action, understand the results (genes and pathways) in the context of drug treatment of UC, CD and psoriasis. For example, to obtain remission related genes for UC, the query was *“list genes that are involved in remission of ulcerative colitis after treatment particularly involved in mucosal or epithelium healing. list it as a python dictionary with gene as key and evidence as value*.*”* Given the remission genes are related to the the type of treatment, the results were manually checked by a immune disease expert for accuracy. The LLMs were also used for initial code generation tasks and researching various GNN architectures.

## Supplementary information

Supplementary information is included in the supplementary_info.pdf file.

## Acknowledgments

We would like to thank Yohan Pierre Lefol from University of Oslo for helping with using TiSA in a temporal setting.

## 6 Declarations

## Funding

Funding was provided by Sanofi

## Conflict of interest

Sachin Mathur, Peyman Passaban, and Hamid Mattoo are Sanofi employees and may hold shares and/or stock options in the company.

## Data availability

All data used for the study is available publicly and in the code repository.

## Code availability

Code is available in a public GitHub repository - https://github.com/Sanofi-Public/GNN-Timeseries

## Author contribution

SM conceived the study, developed methodology, developed code, contributed to functional analysis, and wrote the manuscript. AK was involved in discussions and developed the initial code. AK’s work was done during his internship at Sanofi. PP was involved in discussions, reviewed and edited the manuscript. HM conducted functional analysis, reviewed and edited the manuscript. EH was involved in discussions, gave pivotal ideas, and reviewed the manuscript. ZBJ provided guidance on methods, edited and reviewed the manuscript. Most of the work was done when ZBJ was employed at Sanofi. All authors reviewed the article and approved it for publication.

## Notes

### Competing Interest Statement

The authors have declared no competing interest.

